# Identification and characterisation of potent anti-MRSA phloroglucinol derivatives of *Dryopteris crassirhizoma* Nakai

**DOI:** 10.1101/2022.05.23.493049

**Authors:** Sumana Bhowmick, Manfred Beckmann, Jianying Shen, Luis A.J. Mur

## Abstract

Traditional Chinese Medicine (TCM) has been used to treat infectious diseases and could offer potential drug leads. This study evaluates the in vitro antimicrobial activities commercially sourced *Dryopteris crassirhizoma* Nakai whose authenticity was confirmed by DNA barcoding based on the ribulose bisphosphate carboxylase (*rbcL*) gene. Powdered rhizomes were sequentially extracted using *n-*hexane, dichloromethane, ethyl acetate and methanol at ambient temperature. The dried extracts at different concentrations were tested for antimicrobial activities against *Escherichia coli, Pseudomonas aeruginosa, Staphylococcus aureus*, Methicillin-resistant *Staphylococcus aureus* (MRSA), and *Mycobacterium smegmatis. D. crassirhizoma* extracts exhibited significant antimicrobial activities only against MRSA. Activity-led fractionations of *D. crassirhizoma* and characterisation by Ultra performance liquid chromatography - tandem mass spectrometry (UPLC-MS/MS) identified two potent anti-MRSA phloroglucinol derivatives: Norflavaspidic acid AB and flavaspidic acid AB. The impact of norflavaspidic acid AB on MRSA cells was examined using untargeted metabolomic analysis and compared to that of other established antibiotics (all treatments normalized to MIC_50_ at 6 h). This suggested that norflavaspidic acid AB had a distinctive effect which involved targeting bioenergetic transformation, metabolism, and particularly acetyl CoA, in MRSA cells. No cytotoxicity was observed for norflavaspidic acid AB against murine HepG2 cells. This study requires further experimental validation but can have indicated a naturally available compound that could help counter the threat of clinically relevant strains with antibiotic resistance.

## 1. Introduction

The diversity within the plant kingdom leads it to being the source of highly diverse bioactive compounds. Likely sources of bioactives can be suggested from Ayurveda (Traditional Indian Medicine) dates from about 1000 BC and Traditional Chinese Medicine (TCM) from1100 BC. The earliest records of the uses of a natural product is from 2600 BC which describes the use of oils from *Cupressus sempervirens* (Cypress), *Commiphora species* (myrrh), *Glycyrrhiza glabra* (licorice) and *Papaver somniferum* (poppy juice) to treat cough, colds, parasitic infections and inflammation (Cragg & Newman, 2005). Examples of how such ancients medical systems can serve as sources of what, ultimately, have undergone development and have been marketed as drugs include Paclitaxel (Taxol®) from *Taxus brevifolia* for lung, ovarian and breast cancer (Kaufman et al., 2010), Artemisinin from traditional Chinese plant *Artemisia annua* to combat multidrug resistant malaria (Cragg & Newman, 2005; Paul, 2002), Silymarin extracted from the seeds of *Silybum marianum* for the treatment of liver diseases (Shaker et al., 2010). TCM includes over 7000 plant species have medicinal uses and is attracting global interest as a source of natural products. However, information regarding their use is often inaccessible to the wider scientific community, and there remain difficulties in targeting the most appropriate sources, and product quality assessments.

In terms of a target for TCM-derived drugs, perhaps helping to address the challenge of antimicrobial resistance (AMR) is one of the most pressing. The rise of AMR is recognized as one of the greatest threats to human health with ∼1 million deaths linked to drug resistant microbes between 2014 - 2016 (O’neill, 2016). This reflects that, after decades marked by the discovery of such as sulphonamides, β-lactams, aminoglycoside and trimethoprins, the period after 1980 is defined as an “antibiotic discovery void” (Silver, 2011). Three major factors determine underlie the rise of clinically relevant AMR organisms; (1) the widespread use of antimicrobials; (2) the large and globally connected human population allowing bacterial spread and (3) the extensive and often unnecessary use of antimicrobials providing a strong selective pressure (Michael et al., 2014). One dangerous example of AMR bacterial strains is represented by methicillin-resistant *Staphylococcus aureus* (MRSA). MRSA is a major nosocomial pathogen that has emerged from a hospital setting. At one stage, two strains types, (USA 300 and USA 400) were the primary causes of community-acquired MRSA infections in the USA (McDougal et al., 2003). The MRSA strains are further differentiated based on two types of community-acquired MRSA (CA-MRSA) and hospital-acquired MRSA (HA-MRSA). In general, HA-MRSA strains exhibit high-level resistance to multiple non-β-lactam antimicrobial agents such as quinolones, aminoglycosides, and macrolides. In contrast, CA-MRSA strains are usually susceptible to non-β-lactams, but produce various virulence and colonization factors (Diep & Otto, 2008; Dinges et al., 2000; Zetola et al., 2005). Currently, treatments for severe MRSA infections are vancomycin and daptomycin for bacteraemia; vancomycin, daptomycin or linezolid for complicated skin and soft-tissue infections; and vancomycin or linezolid for hospital-associated pneumonia. However, the overuse of vancomycin in hospitals have resulted in vancomycin resistant MRSA. Linezolid is still used intensively as currently resistance to linezolid remains rare (Itani et al., 2010). Daptomycin also remain effective, but its cost is limiting it use (Bounthavong et al., 2011).

*Dryopteris crassirhizoma* Nakai, a perennial herbaceous fern, known as “the king of antivirals”, is widely distributed in Korea, China, and Japan (Gao et al., 2008). The roots, known as “Gwanjung” in Korea; “Guan Zhong” in China and “Oshida” in Japan, are used in TCM to treat parasitic infestation, haemorrhage, epidemic flu, cold, and cancer (Committee, 1999; “Encycl. Ref. Tradit. Chinese Med.,” 2003; Shinozaki et al., 2008). Powdered and dried rhizomes of various *Dryopteris* ferns have been used as remedies for helminthiasis caused by *Diphyllobothrium latum* (Murakami & Tanaka, 1988). Previously reported phytochemical constituents include triterpene, phloroglucinol, flavonoids and other phenolic compounds(Chang et al., 2006; Min et al., 1998; Noro et al., 1973; Shiojima et al., 1990).

Herein, we screen *Dryopteris crassirhizoma* for antibacterial activities. The most potent activities were identified as phloroglucinol derivatives whose mechanism of action was suggested by metabolomic studies to be linked to a perturbation of glycolysis in MRSA which appeared to significantly deplete acetyl Coenzyme A (acetyl CoA).

## 2. Results

### 2.1. Bioassay-guided isolation bioactives from D. crassirhizoma

DNA barcoding based on the *rbcL* (ribulose-bisphosphate carboxylase) gene was used to authenticate *D. crassirhizoma* as it obtained from commercial sources as dried samples. The sequences (accession number: MN431197) were compared to NCBI voucher sequences to confirm the identity of the TCM.

Powdered samples were sequentially extracted using *n-*hexane, dichloromethane (DCM), ethyl acetate (EtOAc) and methanol (MeOH). Minimal inhibitory concentrations (MIC) of each extract were then screened against a range of clinically relevant bacterial strains, *E. coli* (E), *P. aeruginosa* (P), *S. aureus* (S), MRSA, and *M. smegmatis* (M) (Table 1). The *n-*hexane extract of *D. crassirhizoma* was active against S (MIC of 3.125 µg/mL) and against MRSA (MIC of 3.125 µg/mL). The DCM extract was active against S (MIC of 25 µg/mL) and against MRSA (MIC of 50 µg/mL).

**Table 1:** Antimicrobial screening of crude *D. crassirhizoma* for its minimum inhibitory concentrations [MIC] µg/mL.

| Plant species | Accession number | Part | Solvents | E. | P | M | S | MRSA |
| --- | --- | --- | --- | --- | --- | --- | --- | --- |
| *Dryopteris crassirhizoma* | MN431197 | Rhizome | *n*-Hexane | >500 | >500 | 500 | 3.125 | 3.125 |
|  |  |  | DCM | >500 | >500 | 500 | 25 | 50 |
|  |  |  | EtOAc | >500 | >500 | >500 | >500 | >500 |
|  |  |  | MeOH | >500 | >500 | >500 | >500 | >500 |

E = *Escherichia coli* ATCC 25922, P= *Pseudomonas aeruginosa* ATCC 27853, *Mycobacterium smegmatis* NCTC 333, S = *Staphylococcus aureus* ATCC 29213, MRSA = methicillin-resistant *Staphylococcus aureus* USA300.

Based on the MIC against MRSA observed from crude extracts, we focused on the *n-*hexane extract of *D. crassirhizoma* for further purification (Figure 1). After a first round of purification, assays of the resulting fraction indicated HB displayed the highest anti-microbial activity (MIC of 3.125 μg/mL). Fraction HB was further purified and assessed and fraction of HB5 and HB6 (MIC of 3.125 μg/mL respectively) had the highest activity against MRSA. HB5 was used for further purification based on its higher quantity and the similarity of its mass ions (*m/z*) with HB6 following UHPLC-MS (Figure S1A). Fractions HB5d and HB5e were chosen to be further purified based on their MICs. However, the fractions showed to have very similar biochemical components as detected by UHPLC-MS (Figure S1B) and so were combined and further fractionated. HB5d/e3 (MIC, 6.25 µg/mL) was the most active fraction and UHPLC-MS indicated the abundance of *m/z* 405.15417 with a minor level of *m/z* 419.1695 in positive ionisation mode (Figure S2). HB5d/e5 (MIC, 12.5 µg/mL), also contained only *m/z* 405.15417 and *m/z* 419.1695 but the intensities were more equal. This implied that *m/z* 405.15417 represented the more potent anti-microbial. The fractions underwent several alternative purification techniques, but it was not possible to separate the compounds.

**Figure 1:**
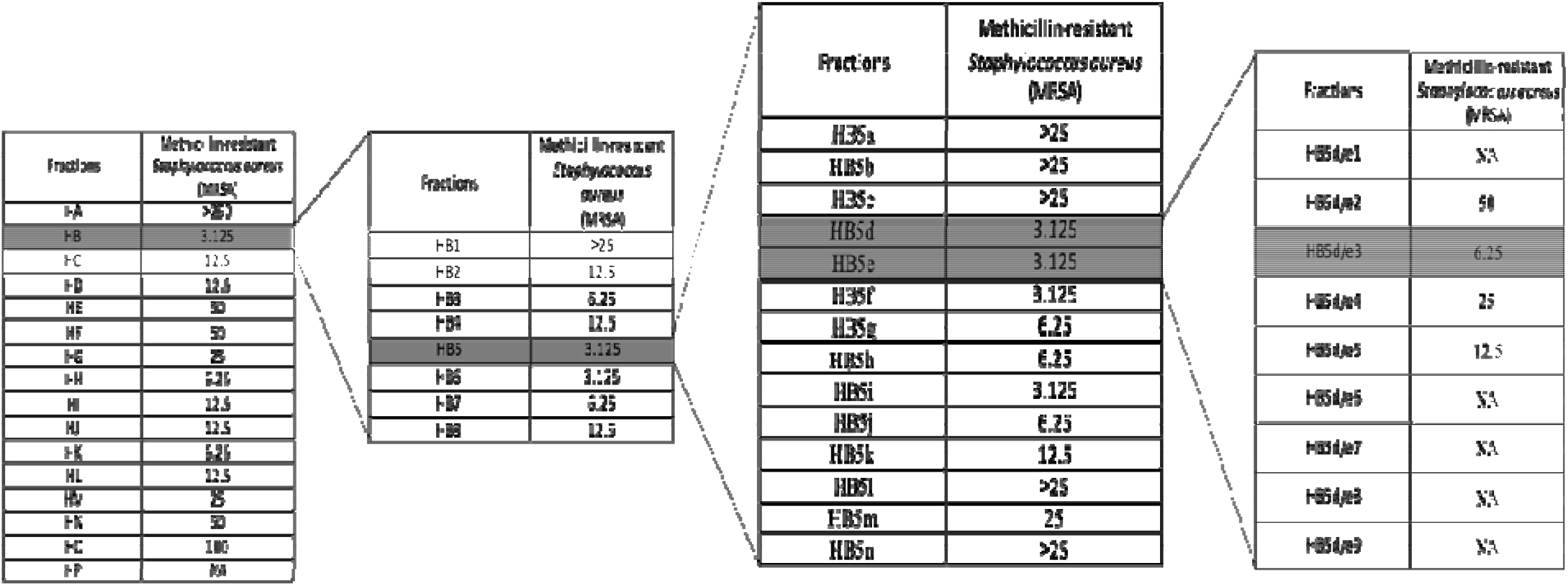
Bioassay guided purified fractions of n-hexane extracts (MIC µg/mL). Note: NA: Not active The components in HB5d/e3 were then characterized and identified via UHPLC-MS coupled with MS^2^/MS^3^ fragmentation based on retention time, accurate mass, elemental composition, and multiple-stage mass data (Table 2, Figures S3 and S4). This suggested that the two compounds were phloroglucinol derivatives, and when compare with the literature, these were identified as norflavaspidic acid AB and flavaspidic acid AB (Ren et al., 2016) (Figure 2).

Given the potent anti-microbial activity of the phloroglucinol derivatives, the cytotoxicity of HB5d/e3 (for simplicity, designed A3) was assessed in HepG2 cell cultures. Screening indicated that the fractions did not show any cytotoxic activities CC50 > 100 µM (> 40.4 µg/mL). The selectivity index (SI) (CC50 / MIC) of the A3 for HepG2 evaluated in relation to their very potent antibacterial activity against MRSA was > 13, which was favourable for further development.

**Table 2:**
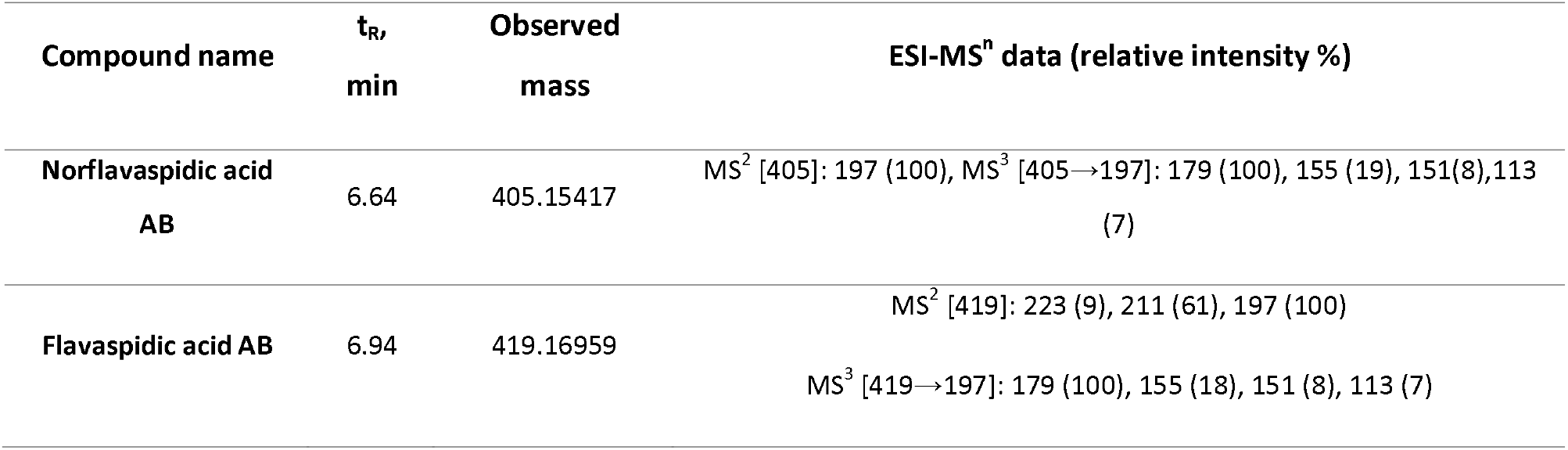
Identification of antimicrobial metabolites from D. crassirhizoma using UHPLC-MS/MS

### 2.2. Untargeted metabolomics suggested the activities of the phloroglucinol derivatives

To suggest the defining possible mode of action for A3 on MRSA, a metabolomic was employed. This involved defining the concentrations of A3 and a range of established antibiotics with different known cellular targets required to inhibit MRSA growth by 50 % at 6 h (MIC_50_) (Table 3, Figure S5).

**Table 3:**
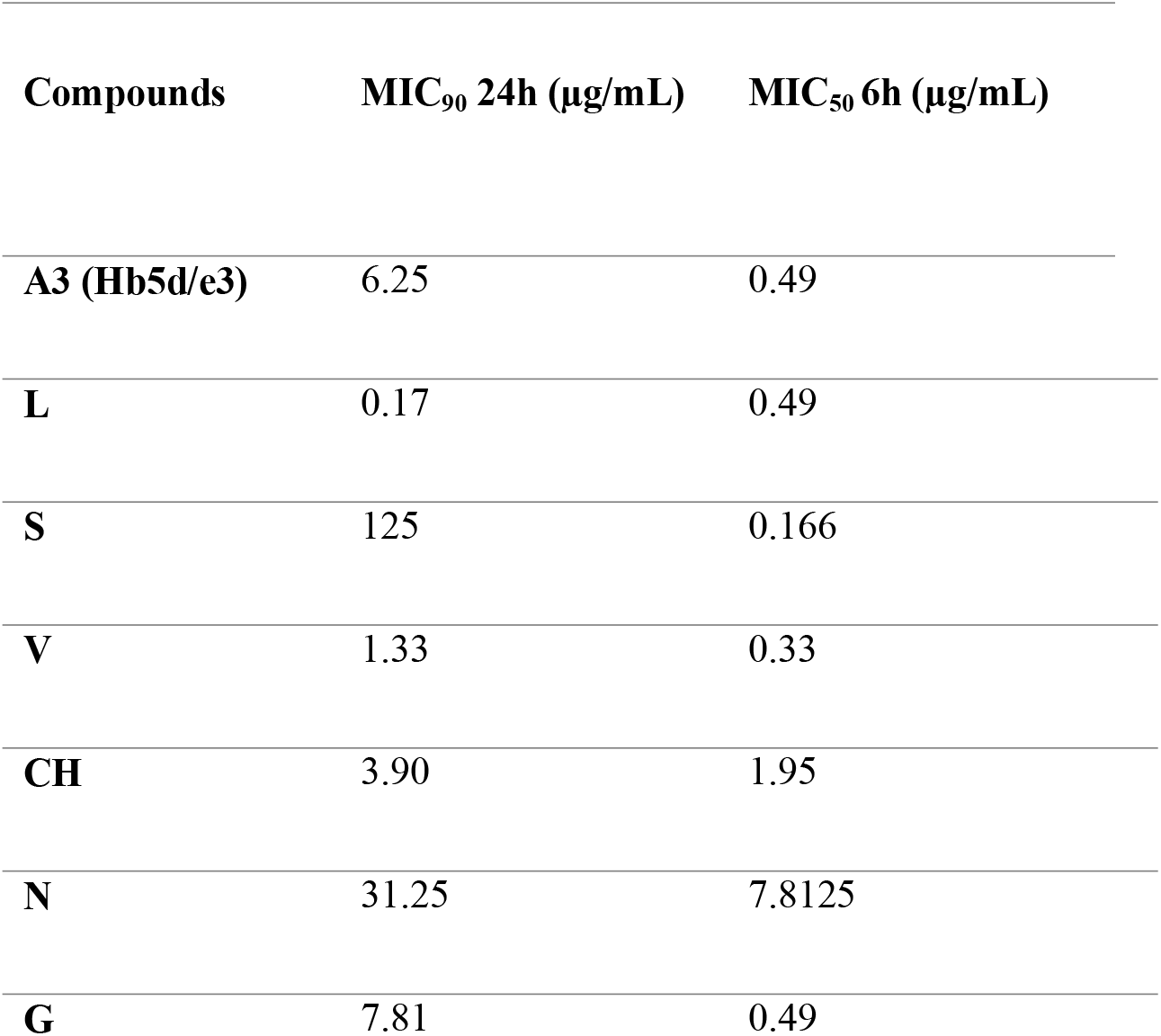
Standardisation of antibiotics based on their MICs.

The concentration of antibiotic required to inhibit growth of USA300 MRSA at 24 h and by 50 % over 6 h at a bacterial concentration of 1×10^8^ CFU/mL. Note: A3 (Hb5d/e3), chloramphenicol (CH), gentamycin (G), levofloxacin (L), nalidixic acid (N), streptomycin (S), vancomycin (V).

Flow infusion electrospray ionization high resolution mass spectrometry (FIE-HRMS) was used to profile extracted metabolites derived from MRSA treated with A3 versus established antibiotics chloramphenicol (CH), gentamycin (G), levofloxacin (L), nalidixic acid (N), streptomycin (S), vancomycin (V) and control MRSA cells (Figure 3). In Figure 3, unsupervised PCA considered the established antibiotics based on their major modes of action. This indicated that immediately after treatment, the MRSA metabolomes were identical (Figure 3A). However, within 2 h of treatment with A3 resulted in a shift in metabolome that was distinct from the responses to established antibiotics. It was notable that the metabolomes of the established antibiotics clustered together and were therefore fundamentally similar, irrespective of their different modes of action. This did not arise from the MRSA cells being all dead, as the concentrations of antibiotic only suppressed bacterial growth by 50 % at 6 h. However, this could indicate that the MRSA metabolomes were reflecting a common stress response by the bacteria

The *m/z* that differed significantly (*P=* 0.001) between control and A3 treated were identified by ANOVA corrected for FDR. These were identified using the “MS peaks to pathway” function on MetaboAnalyst 4.0 (Li et al., 2013) and mapped on the KEGG general metabolism map (Figure 4). It was notable that the vast majority of the significantly affected pathway involved glycolysis. To provide further insights into the effects of A3 (Hb5d/e3) on MRSA metabolism, the significantly differing metabolites were displayed on to heatmap (Figure 5). This indicated that the levels of glycolysis intermediates-such as phosphoenolpyruvate (PEP) and 3-phosphoglycerate (3-PG)-were greatly decreased in cells treated with A3 (Figure S6). This shift in the glycolysis/gluconeogenesis pathway did not arise from a shift towards the production of lactate or to the citrate cycle since these were not targeted as being affected in our assessments. However, the dramatic loss in acetyl CoA observed can undoubtedly affect the production of citrate and therefore the function of the TCA cycle.

## 3. Discussion

The rise of AMR and the discovery void in the derivation of novel antibiotics has led a revisiting of traditional medicine as sources of natural products which could represent drug leads. China has been using TCM herbs for centuries and these are now attracting global interest as a source of natural products. However, the information regarding their use is often inaccessible to the wider scientific community. In this collaborative study, we engaged in a screen of potential antimicrobial properties and the targeting of antimicrobial metabolites from *D. crassirhizoma*.

Based on our studies, we suggest that the quality, precise identification, and reliability in the plant species from where the natural product is obtained is very critical step for successful innovative drug discovery. The complex chemical composition of plants or TCM material and variation in source can lead to batch-to-batch inconsistency. Genomic techniques such as DNA barcoding are established techniques that rely on sequence diversity in short, standard *rbcL* regions (400–800 bp) and internal transcribed spacer for species-level identification(Ganie et al., 2015). Thus, we have incorporated a quality assessment based on DNA barcoding.

*D. crassirhizoma* exhibited particularly potent activity against MRSA. Sequential bioactivity guided fractionations lead the identification of two phloroglucinols metabolites: Norflavaspidic acid AB and flavaspidic acid AB. The close similarities in their chemical structures (Figure 2) prevented them being separated with the use of preparative HPLC or TLC. However, the considerable predominance of norflavaspidic acid AB in the most active fraction, implies that this had the greater anti-MRSA properties compared to flavaspidic acid AB. Norflavaspidic acid AB is an example of acyl phloroglucinol dimers formed by a methylene linkage. In terms of reported bioactivities, no antimicrobial activity has been previously reported but norflavaspidic acid AB has inhibitory effects on melanin production by melanoma B16F10 cells with IC50 values of 181.3 μM (Pham et al., 2017) and against human leukemia Reh cells with IC50 of 32.2 µg/mL (Ren et al., 2016). Further, it countered the effects of the pro-apoptotic factor - FAS - of IC50 29.7µM (Na et al., 2006). However, we observed no cytotoxic effects for A3 against HepG2 cells where norflavaspidic acid AB was predominant. Flavaspidic acid AB has been reported to have antibacterial activity based on when a paper disc diffusion assays with MICs ranging between 12-20 µg/mL depending on the microorganism being tested (H. B. Lee et al., 2009) which is in line with our observations. Moreover, flavaspidic acid AB exhibited a potent antioxidant activity against the LPO inhibitory test with IC50 values of 13.1 mM (S. M. Lee et al., 2003). It can induce IFN-α, IFN-β, and IL1-β expression in porcine alveolar macrophages, which could contribute to inhibition of porcine reproductive and respiratory syndrome virus (PRRSV) replication (Yang et al., 2013) It also shows inhibitory effects against human leukemia Reh cells with IC50 of 35.3 µg/mL (Ren et al., 2016). Its activity against the pro-apoptotic ligand FAS had an IC50 of 28.7µM (Na et al., 2006). It is now important to derive pure norflavaspidic acid AB and flavaspidic acid AB so that the anti-microbial properties of each can be properly assessed. This could be important given that we could find no evidence of cytotoxicity with these metabolites and the potencies against a clinically relevant MRSA strain.

**Figure 2:**
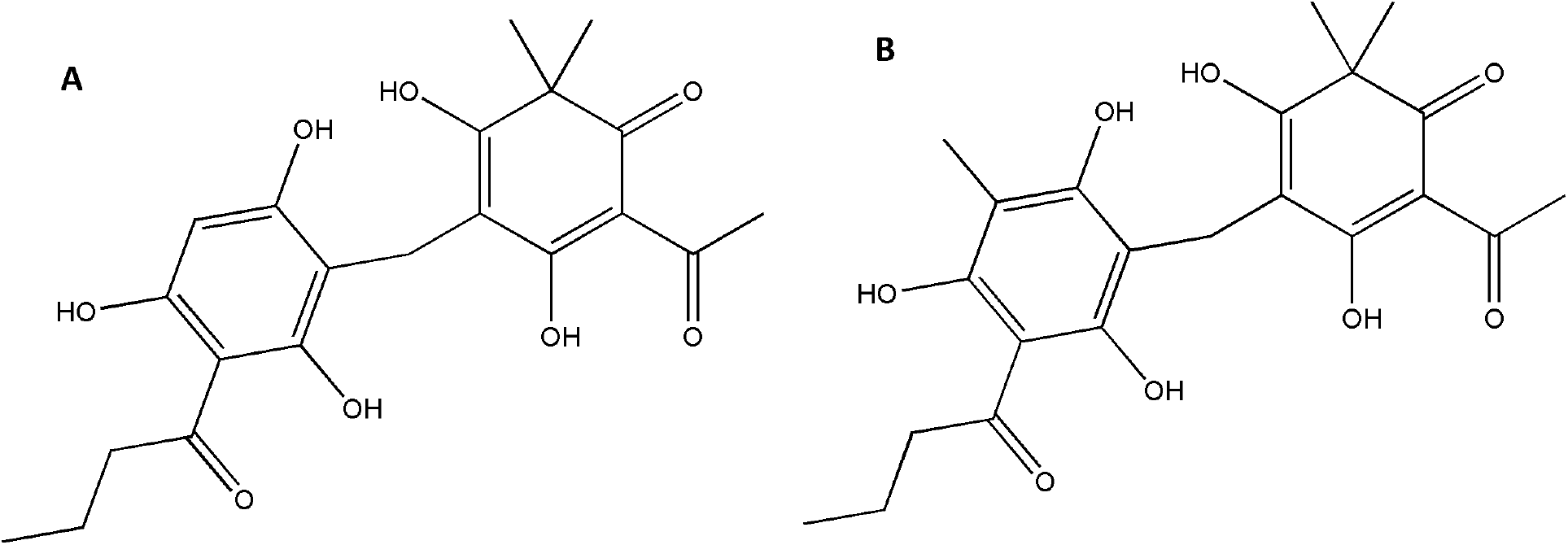
The structures of the bioactive natural products isolated from *Drypotersis crassirhizoma*: (**A**) Norflavaspidic acid AB and (**B**) Flavaspidic acid

**Figure 3:**
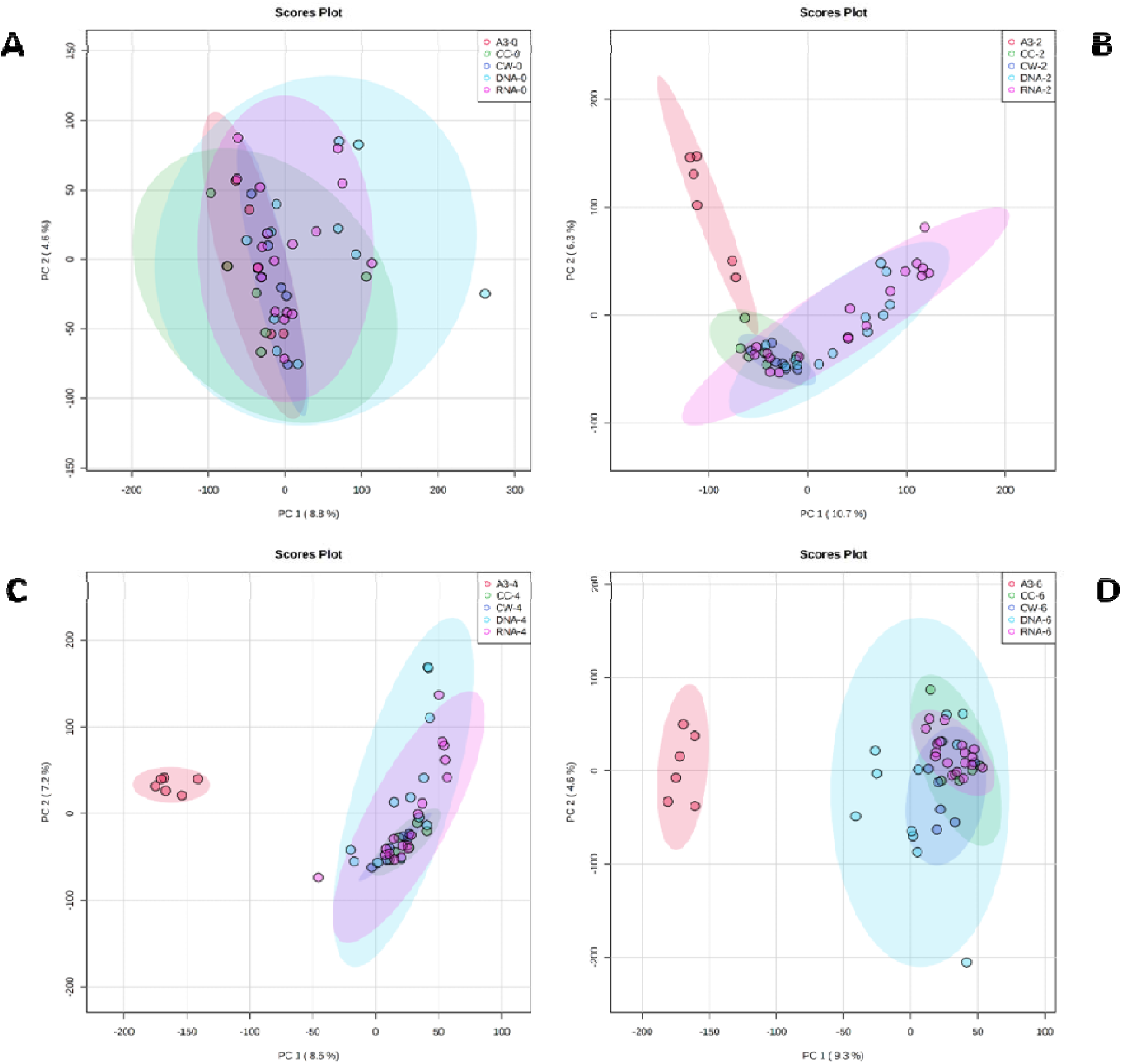
Principal component analysis (PCA) of antibiotic treated MRSA metabolomes. PCA score plots (95 % confidence interval illustrated show as paler coloured ellipses) of normalized m/z intensities of metabolites extracted as detected by flow infusion electrospray high-resolution mass spectrometry (FIE-HRMS).from MRSA treated with A3 (Hb5d/e3) and compared to control bacteria (CC) and to bacteria treated with antibiotics with similar mechanism of action, grouped into those with activity on cell wall (vancomycin, CW), on DNA (Levofloxacin and Nalidixic acid) and on RNA (Chloramphenicol, Gentamycin and Streptomycin) for bot positive and negative mode for time points (A) 0, (B) 2, (C) 4 and (D) 6 h.

**Figure 4:**
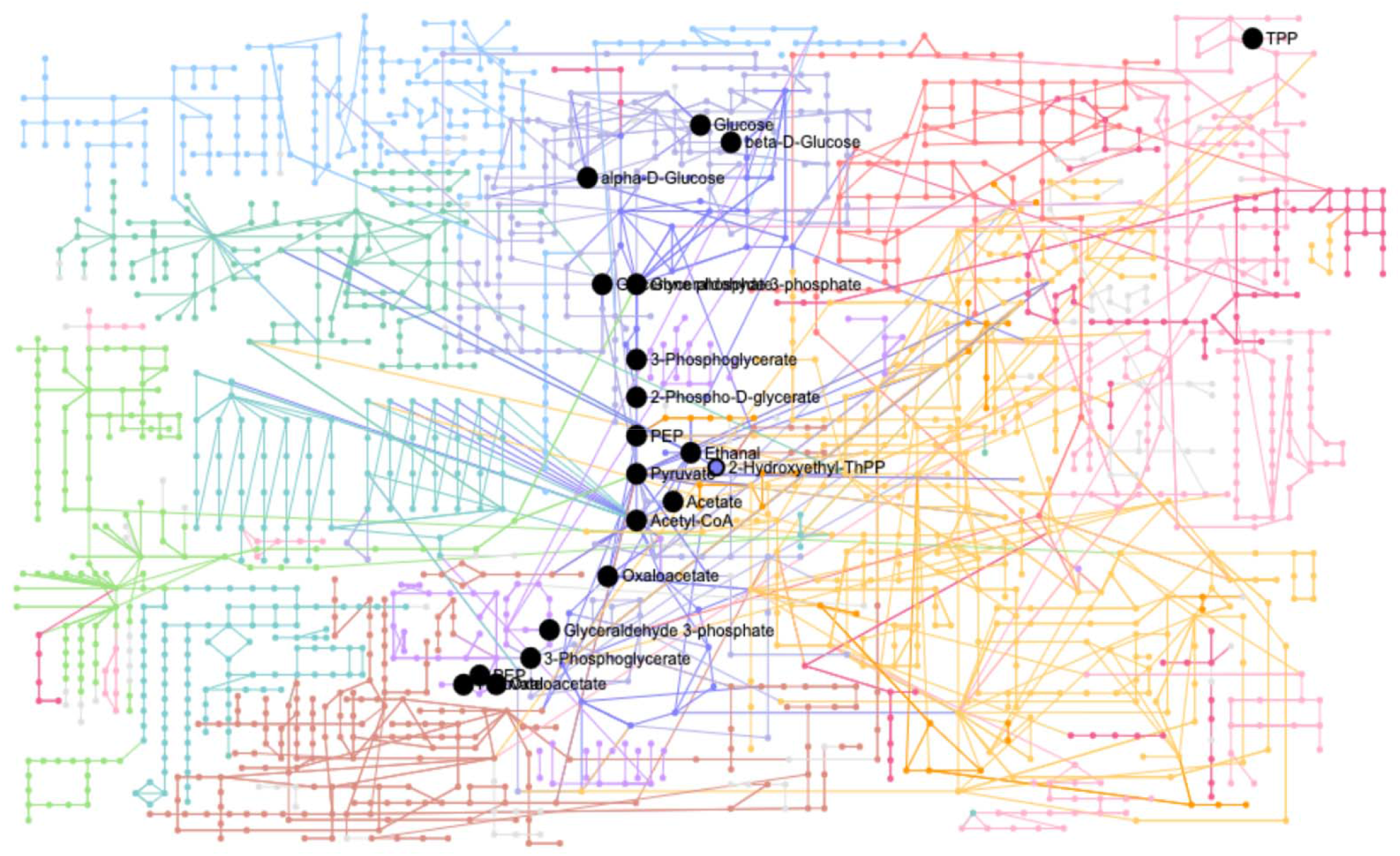
Mapping significant metabolite changes of MRSA in both treated A3 (Hb5d/e3) and non-treated cells (CC) on to cellular metabolism as defined by KEGG

**Figure 5:**
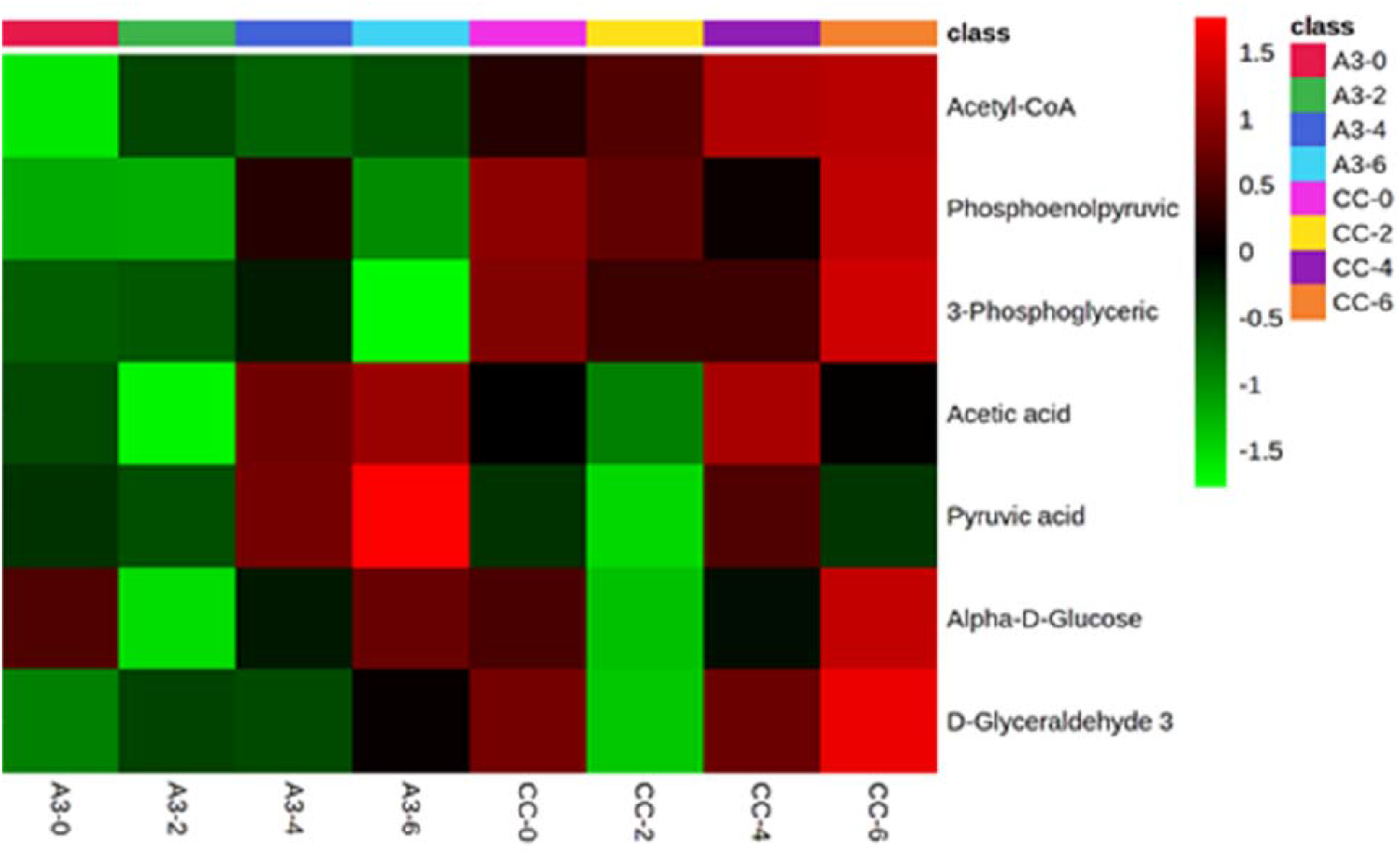
Heatmap of all the significantly different metabolite changes of glycolysis and gluconeogenesis pathway happening at different time points in both treated A3 (Hb5d/e3) and non-treated cells (CC).

To provide more information on how norflavaspidic acid AB/ flavaspidic acid AB acts we used metabolomic assessments in comparison to the established antibiotics based on their different modes of action. There are a series of known anti-microbial target in MRSA these include cell wall inhibitors. Anti-MRSA β-lactam antibiotics include ceftobiprole and ceftaroline. Ceftobiprole is the first molecule from a new class of cephalosporins that was developed to bind specifically to mutant Penicillin-Binding Proteins (PBPs) in MRSA (Hamad, 2010) and resists the action of class A (TEM-1) and class C (AmpC) β-lactamases(Queenan et al., 2007) and is presently in Phase 3 clinical trials (Bassetti & Righi, 2015). However, rapid emergence of methicillin-resistant *S. aureus* (MRSA) containing modified penicillin-binding protein (PBP) 2a encoding operon was poorly inhibited by β-lactams (Grundmann et al., 2006). This is overcome by the glycopeptide vancomycin and has been the drug of choice for MRSA treatment for numerous years. Vancomycin acts by inhibiting proper cell wall synthesis but almost exclusively on in Gram-positive bacteria. However, because of limited tissue distribution, as well as the emergence of isolates with reduced susceptibility and in vitro resistance to vancomycin, the need for alternative therapies that target MRSA has become apparent(Bassetti & Righi, 2015; Hidayat et al., 2006; Micek, 2007). Alternatives can include aminoglycosides – for example, gentamycin - that block bacterial protein synthesis by inducing codon misreading and interrupting translocation of the complex between tRNA and mRNA, interacting directly with the 30S rRNA(Carter et al., 2000). Other options are DNA gyrase and topoisomerase inhibitors which include Levofloxacin and Nalidixic acid. New generation quinolones such as delafloxacin and finafloxacin show excellent results against *S. aureus* strains and specifically, MRSA (Kocsis et al., 2021).

Our metabolomic comparison with A3 included examples of different forms of anti-MRSA activities. These appeared to have similar effects at 6 h following treatment at concentrations sufficient to suppress MRSA growth by 50%. Given the different modes of action this suggestive of revealing a common effect such as the slowing of growth and/ or associated bacterial stress. However, the A3 effect was strikingly different suggesting a different mechanism of action to the other established antibiotics. When identifying the key metabolomic differences with A3 treatment, these appeared to be linked to altered bioenergetic mechanisms, particularly associated with the perturbation of glycolysis. Although our assessments could not identify the exact targets, for example, an enzyme, that could be being inhibited by A3, the consequences appeared to lead to a collapse in the levels of acetyl CoA. Acetyl CoA due to its importance in metabolism it has been suggested to be one of a series of key “sentinel” metabolites that indicates the metabolic state of the cell(Shi & Tu, 2015). The targeting of glycolysis by A3 would explain the loss in acetyl CoA as this provides the pyruvate which can be oxidative decarboxylated to produce acetyl CoA. In mammalian systems the depletion of acetyl CoA can lead to autophagy (Mariño et al., 2014) and in *Mycobacterium tuberculosis* perturbation of acetyl CoA metabolism was clearly indicated to be a killing mechanism (Beites et al., 2021). Following is condensation with oxaloacetate it can form citrate which can the feed into the TCA cycle, and it was surprising that this was not apparently significantly altered in our study, although TCA intermediates are relatively easy to detect using our metabolomic technology. Interesting, flavaspidic acids have been suggested to affect cellular respiration and oxidative phosphorylation in isolated hepatocytes and even, at moderate concentrations, uncouples oxidative phosphorylation in isolated hepatocytes (Burnett et al., 1979). This would align with our observations of anti-MRSA activities, but such a cross-kingdom effect would not agree with the lack of cytotoxicity against HepG2 cells.

## 4. Materials and Methods

### 4.1 Plant materials

The commercial sourced dried plant material was provided by Artemisinin Research Centre, Institute of Chinese Materia Medica, China Academy of Chinese Medical Sciences, Beijing, China.

### 4.2 DNA barcoding of plant samples

Total genomic DNA of was extracted using a DNeasy Plant Mini kit (Qiagen, UK) in accordance with the manufacturer’s instructions. The *rbcL* sequences where amplified with *rbcL*a-F: 5′-ATGTCACCACAAACAGAGACTAAAGC-3′ and *rbcL*a-R: 5′-GTAAAATCAAGTCCACCRCG-3′; DNA barcoding primers (Hollingsworth, 2011). PCR used a total volume of 20 µL with 10 µL BioMix (BioLine, UK) 1 µL of each primer (10 µM), and 1 µL of the DNA template (50 ng/ µL) and 7 µL distilled water. PCR involved 1 cycle (94 °C for 3 min), 35 cycles (94 °C for 1 min, 55 °C for 1 min, and 72 °C for 1 min), and 1 cycle 72 °C for 7 min. The resulting 600 bp band was sequenced 3730xl DNA Analyzer (Applied Biosystems, UK), using the same primers as used for PCR. The derived sequences were assessed using BLAST (https://blast.ncbi.nlm.nih.gov/Blast.cgi). The *rbcL* sequence was submitted to Genbank (Accession number: MN431197).

### 4.3 Sequence of bioassay guided purification

800g of dried material of *D. crassirhizoma* was ground into a powder and was extracted sequentially using *n*-hexane, dichloromethane (DCM), and methanol (MeOH) at room temperature with continuous stirring for 24 h respectively. Bioactivity-linked fractionation of the *n*-hexane extract was based on the ability to suppress the growth MRSA

Approximately 94 g of *n-*hexane extract of *D. crassirhizoma* was fractionated using silica gel column chromatography (CC) (4×50 cm, 150 g of SiO2,). The column eluted with *n-*hexane-EtOAc (1:0 to 0:1, at 10 % gradient, 500×4 mL of each eluent), and EtOAc-MeOH (1:0 to 1:2, 500 mL X 3 of each eluent) to yield 16 fractions (designated H for “*n*-hexane” and A through to P; thus, HA through to HP). Fraction HB (59.11 g) was again subjected to silica CC (4×50 cm, 150 g of SiO2,) eluting with *n-*hexane-EtOAc (1:0 to 3:20, 1 % gradient, 500 mL X 4 of each eluent) and EtOAc-MeOH (1:0 to 1:2, 5 % gradient, 500 mLX5 of each eluent) to yield 8 fractions (1 through 8; thus HB1-HB8). The HB5 fraction (10g) fractionated again using silica gel CC (4 × 50 cm, 150 g of SiO2,) eluted with *n-*hexane-EtOAc (1:0 to 1:10, 1 % gradient, 500 m X 4 of each eluent); *n-*hexane-EtOAc (1:10 to 0:1, 10 % gradient, 500 mL X4 of each eluent) and EtOAc-MeOH (1:0 to 1:2, 10 % gradient, 500 mL X2 of each eluent). This resulted in 14 fractions, a through n: HB5a-HB5n.

Fractions HB5d and HB5e showed similarity in components detected by UHPLC-MS and were combined (800 mg) and further fractionated with silica CC (4×33 cm, 150 g of SiO2). Eluting with *n-*hexane-DCM (1:0 to 0:1, 10 % gradient, 500 mL× 4 of each eluent) and DCM-MeOH (1:0 to 1:2, 500 mL × 4 of each eluent) yielding four fractions HB5d/e1 - HB5d/e8. All the fractions were assessed by thin layer chromatography (TLC) to check for purity.

### 4.4 Bacterial growth and Minimum Inhibitory Concentration (MIC) calculation

All procedures were performed in a biosafety level 2 (BSL2) cabinet. Samples were dissolved in methanol: water (1:1) to give a final concentration of 5 mg/mL. The antimicrobial activity of all the extracts and isolated compounds was evaluated in 96-well plates by micro-dilution technique over a concentration range (500 µg /mL to 1.5625 µg/mL^)^. Serial two-fold dilutions was performed in nutrient broth (for *Escherichia coli* ATCC 25922, *Pseudomonas aeruginosa* ATCC 27853, *S. aureus* ATCC 29213, MRSA = methicillin-resistant *S. aureus* USA300) and nutrient broth supplemented with 0.05 % tween 80 and 0.2 % glycerol (for *Mycobacterium smegmatis* NCTC 333). All diluted extracts and compounds were tested in triplicate in an initial bacterial concentration of 5.0□×□10^5^ CFU/mL. The MIC was determined as the lowest concentration of a drug at which no growth was visible at 37 °C after 24 h for *E. coli, P. aeruginosa, S. aureus*, MRSA and after 72 h with *M. smegmatis*. Rifampicin and gentamycin sulphate (50 µg/mL) were used as positive controls. The optical density OD600 was measured in a Hidex plate (Hidex Sense, UK). Growth was assessed by adding 20 µL 0.5 mg/mL 3-(4, 5-dimethylthiazol-2-yl)-2, 5-diphenyltetrazolium bromide (MTT) stain.

### 4.5 Ultra-high performance liquid chromatography–high resolution mass spectrometry (UHPLC-HRMS)

Fractions were analysed on an Exactive Orbitrap (Thermo Fisher Scientific) mass spectrometer, which was coupled to an Accela Ultra High-Performance Liquid Chromatography (UHPLC) system (Thermo Fisher Scientific). Chromatographic separation was performed on a reverse phase (RP) Hypersil Gold C18 1.9 µm, 2.1□×□150 mm column (Thermo Scientific) using water using 0.1 % formic acid (v/v, pH 2.74) as the mobile phase solvent A and acetonitrile: isopropanol (10:90) with 10 mM ammonium acetate as mobile phase solvent B. Each sample (20 μL) was analysed using 0-20% gradient of B from 0.5 to 1.5 min and then to 100 % in 10.5 min. After 3 min isocratic at 100 % B the column was re-equilibrating with 100 % A for 7 min.

### 4.6 HepG2 Cell Culture and MTT Assay

Human Caucasian hepatocyte carcinoma (HepG2) (sigma Aldrich, cat# 85011430) was used for the cytotoxicity experiments. Targeted fractions were assessed for 50% of cytotoxicity (CC50) at concentrations were from 200 µM to 1 µM in triplicates. Cell viability used the MTT assay as described by (Crusco et al., 2019).

### 4.7 Standardisation of antibiotic dosage

Antibiotics treatments were standardised to concentrations which inhibited growth by 50 % (MIC_50_) at 6 h following microdilution of all antibiotics of a bacterial culture of OD600 0.6 compared to untreated controls. The antibiotics used were chloramphenicol (CH), gentamicin (G), levofloxacin (L), nalidixic acid (N), streptomycin (S) and vancomycin (V), were obtained from Sigma-Aldrich Ltd (UK) and *D. crassirhizoma* metabolites.

### 4.8 Sample preparation for metabolomics

The procedure previously described by Baptista et.al (2018) was followed with minor modifications. The bacteria treated with antibiotic and the untreated control group were independently cultured (n = 6 replicates) at constant shaking at 200 rpm at 37 °C (Gallenkamp Orbital Incubator).

All samples were collected during mid-exponential growth phase. A 3 mL aliquot of bacterial culture (OD600 at time 0□h was 0.6) was harvested at 0, 2, 4 and 6 h after the treatment with each respective antibiotic and controls. The samples were centrifuged at 2 min, 10 °C at 4500 rpm, and the pellet was resuspended with 3 mL of cold saline solution (0.85 % NaCl in H2O w/v).. The samples were stored at −80 °C after the cellular metabolism was rapidly quenched with liquid N2 until analysis. On analysis, samples were thawed and centrifuged at 10 °C at 4500 rpm for 2 min, the pellet washed in 4 mL of cold saline solution (0.85 % NaCl in H2O w/v) and re-centrifuged (10 °C at 4500 rpm for 2 min). All samples were adjusted to an OD600 of 1 and centrifuged to ensure similarly sized bacterial populations were assessed. 70 μL of a chloroform: methanol: wate r (1: 3: 1) mixture was added to the pellet and extractions involved four freeze-thaw cycles with periodic vortexing. After final centrifugation at 4500 rpm, 60 μL of the particle-free supernatant was transferred into a microcentrifuge tube. An additional extraction with 50 μL of chloroform: methanol: water (1: 3: 1) was done and the new supernatant, after final centrifugation, was combined with the supernatant from the first extraction. From this mixture, 50 μL were transferred into an HPLC vial containing a 0.2 mL flat-bottom micro insert for FIE-HRMS analysis.

### 4.9 Metabolomic analyses

Extracted metabolites were analysed by flow infusion electrospray ionization high resolution mass spectrometry (FIE-HRMS) in the High-Resolution Metabolomics Laboratory (HRML), Aberystwyth University. Metabolite fingerprints were created in both positive and negative ionisation mode. Ion intensities were acquired between *m/z* 55 and 1200 for 3.5 min in profiling mode at a resolution setting of 280,000 (at *m/z* 200) resulting in 3 (± 1) ppm mass accuracy. Samples (20□µL volume) were injected by an autosampler into a flow of 100 μL/min methanol/water (70:30, v/v). Electrospray ionization (ESI) source parameters were set based on manufacturer’s recommendations. An in-house data aligning routine in Matlab (R2013b, The MathWorks) was used to join mass spectra around the apex of the infusion maximum into a single mean intensity matrix (runs × *m/z*) for each ionization mode and normalised to total ion count. The derived matrices are shown in Table S1. Data were log10-transformed and used for statistical analysis performed by MetaboAnalyst 4.0 (Chong et al., 2018) which was also used to perform principal component (PCA) and heat maps. The MetaboAnalyst 4.0 - MS peaks to pathways module was used to identify metabolites (tolerance = 3 ppm) and significant affected metabolic pathways (model organism = *S. aureus*) (Chong et al., 2018). Pathway enrichment assessments used the *mummichog* algorithm based on mass spectrometry data, avoiding the *a priori* identification of metabolites (Li et al., 2013). Mummichog plots all possible matches in the metabolic network and then looks for local enrichment, providing reproduction of true activity, as the false matches will distribute randomly (Li et al., 2013).

## 5. Conclusions

The study also attempts in characterizing the bioactives in *D. crassirhizoma* responsible for the anti-MRSA activity. Norflavaspidic acid AB and flavaspidic acid AB was identified as the major components of the fractions. There were several alternative methods used to separate the fraction but failed. The mode of action of A3 was further explored using metabolomics. Although it is currently not unequivocally established how A3 causes the observed

MRSA cell death, our metabolomic approach suggested some potential mechanisms. Whether one or more of these mechanisms are main effects or how far they are to the so-called off-target effects requires further work. However, this study demonstrated the phloroglucinol derivatives in *D crassirhizoma* including norflavaspidic acid AB and flavaspidic acid AB have potent antibacterial activity against MRSA. One or both compounds may be a new class of anti-MRSA antibiotics.

## Supplementary Materials

**Figure S1:** Total ion count (TIC) chromatograms of (A) Total ion count (TIC) chromatograms of HB5 and HB3 obtained by UHPLC-MS.(i) Positive Ions of HB5 (black) and HB6 (red); (ii) Negative ions of HB5 (black) and HB6 (red) (B) HB5d and HB5e obtained by UHPLC-MS. (iii) Positive Ions of HB5d (black) and HB5e (red); (iv) Negative ions of HB5d (black) and HB5e (red).

**Figure S2:** TIC chromatograms of HB5d/e3 (A3) at positive ionization. A: Total ion chromatograms of peak of HB5d/e3 (A3) at positive ionization (base peak intensity:1.86E8). B: Mass range 404.5 - 405.5 was selected (base peak intensity: 3.63E8). C: Mass range 418.5 – 419.5.5 was selected (base peak intensity: 3.82E7).

**Figure S3:** (A) represents the conventional MS/MS spectra data of norflavaspidic acid AB ([MH]+obs. = 405.15413 amu). A peak at *m/z* 419 and 441 were contaminations from the column. (B) Represents the spectrum of the MS/MS/MS experiment in which norflavaspidic acid AB was fragmented by in-source fragmentation and its product ion peak at *m/z* at 405 was selected by Q1 and further fragmented in Q2. A second-generation product ion at *m/z* 197 was a result of breaking of the bond. (C) represents the MS/MS/MS spectrum of the product ion peak at *m/z* 197. A main product ion at m/z 179 accompanied a neutral loss of 18 amu. The spectra also contain other peaks at *m/z* 154(loss of loss of 25 amu), *m/z* 151 (loss of 28 amu), *m/z* 161(loss of 18 amu) and *m/z* 137 (loss of 42 amu).

**Figure S4: (**A) represents the conventional MS/MS spectra data of flavaspidic acid AB ([MH]+obs. = 405.16965 amu). A peak at *m/z* 441 were contaminations from the column. (B) Represents the spectrum of the MS/MS/MS experiment in which flavaspidic acid AB was fragmented by in-source fragmentation and its product ion peak at *m/z* at 405 was selected by Q1 and further fragmented in Q2. A second-generation product ion at *m/z* 211 and 197 was a result of breaking of the bond. (C) represents the MS/MS/MS spectrum of the product ion peak at *m/z* 197. A main product ion at *m/z* 179 accompanied a neutral loss of 18 amu. The spectra also contain other peaks at *m/z* 155(loss of loss of 24 amu), *m/z* 151 (loss of 28 amu), *m/z* 161(loss of 18 amu) and *m/z* 137 (loss of 42 amu);

**Figure S5:** Transmission electron microscope images of MRSA: USA 300 treated with two different concentrations of HB5d/e3 (A3) (A) untreated controls at 6h (B) concentrations of HB5d/e3 (A3) sufficient to inhibit growth by 50 % at 6h (MIC_50_) and (C), and contractions sufficient to inhibit growth by 90 % at MIC90 at 24h. Arrow shows the cell membrane disruption and lysis;

**Figure S6:** Schematic showing metabolite changes in MRSA cell following treatment with A3 based on KEGG map00010 “glycolysis and gluconeogenesis”. Metabolite names which were shown to be reduced with A3 treatment are indicated with a green background. The heatmap showing changes over time in controls (CON) and A3 treatment are also given (also provided in Figure 5). Metabolites or pathways highlighted in grey, showed no significant change in A3 treatment.

**Table S1:** *m/z* matrix derived following flow-infusion electrospray mass spectrometry.

## Author Contributions

Conceptualization, L.M. and J.S.; methodology, S.B. and M.B.; formal analysis, S.B.; resources, M.B.; writing—original draft preparation, S.B.; writing—review and editing, L.M. and J.S.; supervision, L.M. and J.S.; project administration, L.M. and J.S.; funding acquisition, L.M. All authors have read and agreed to the published version of the manuscript.

## Institutional Review Board Statement

Not applicable.

## Informed Consent Statement

Not applicable.

## Data Availability Statement

Not applicable.

## Acknowledgments

We appreciate the technical support provided with Mass Spectroscopy by Helen Phillips (Aberystwyth University, UK). Thanks also to Dr. Gilda Padalino and Prof. Karl Hoffmann (Aberystwyth University, UK) for the cytotoxicity assessments based on HepG2 cells. S.B. was supported by an AberDoc PhD scholarship from Aberystwyth University, UK.

