## Supplementary data for "Identification and characterisation of potent anti-MRSA phloroglucinol derivatives of *Dryopteris crassirhizoma* Nakai"

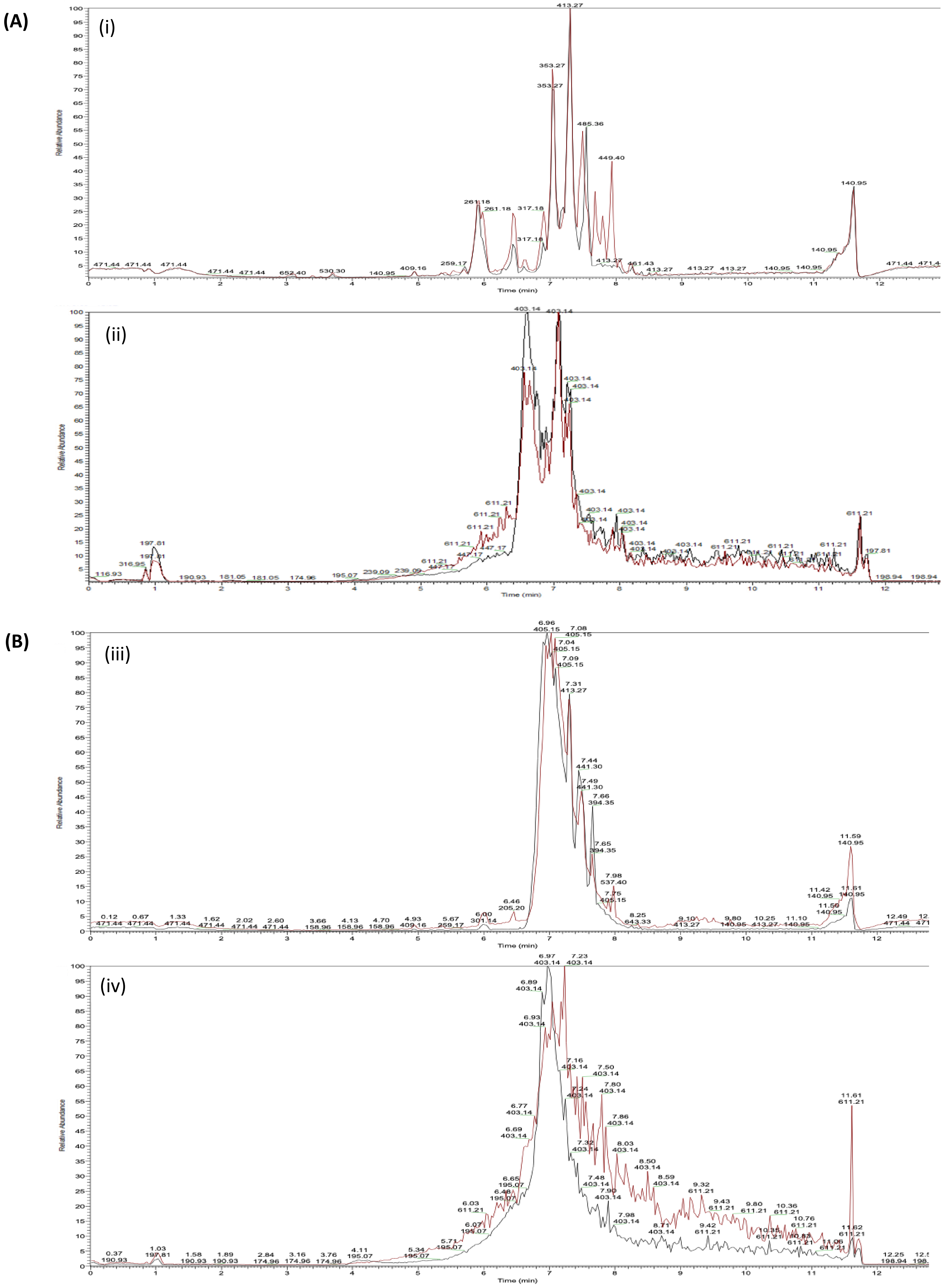


**Supplementary Figure 1**: Total ion count (TIC) chromatograms of (A) Total ion count (TIC) chromatograms of HB5 and HB3 obtained by UHPLC-MS. (i) Positive Ions of HB5 (black) and HB6 (red); (ii) Negative ions of HB5 (black) and HB6 (red) (B) HB5d and HB5e obtained by UHPLC-MS. (iii) Positive Ions of HB5d (black) and HB5e (red); (iv) Negative ions of HB5d (black) and HB5e (red)

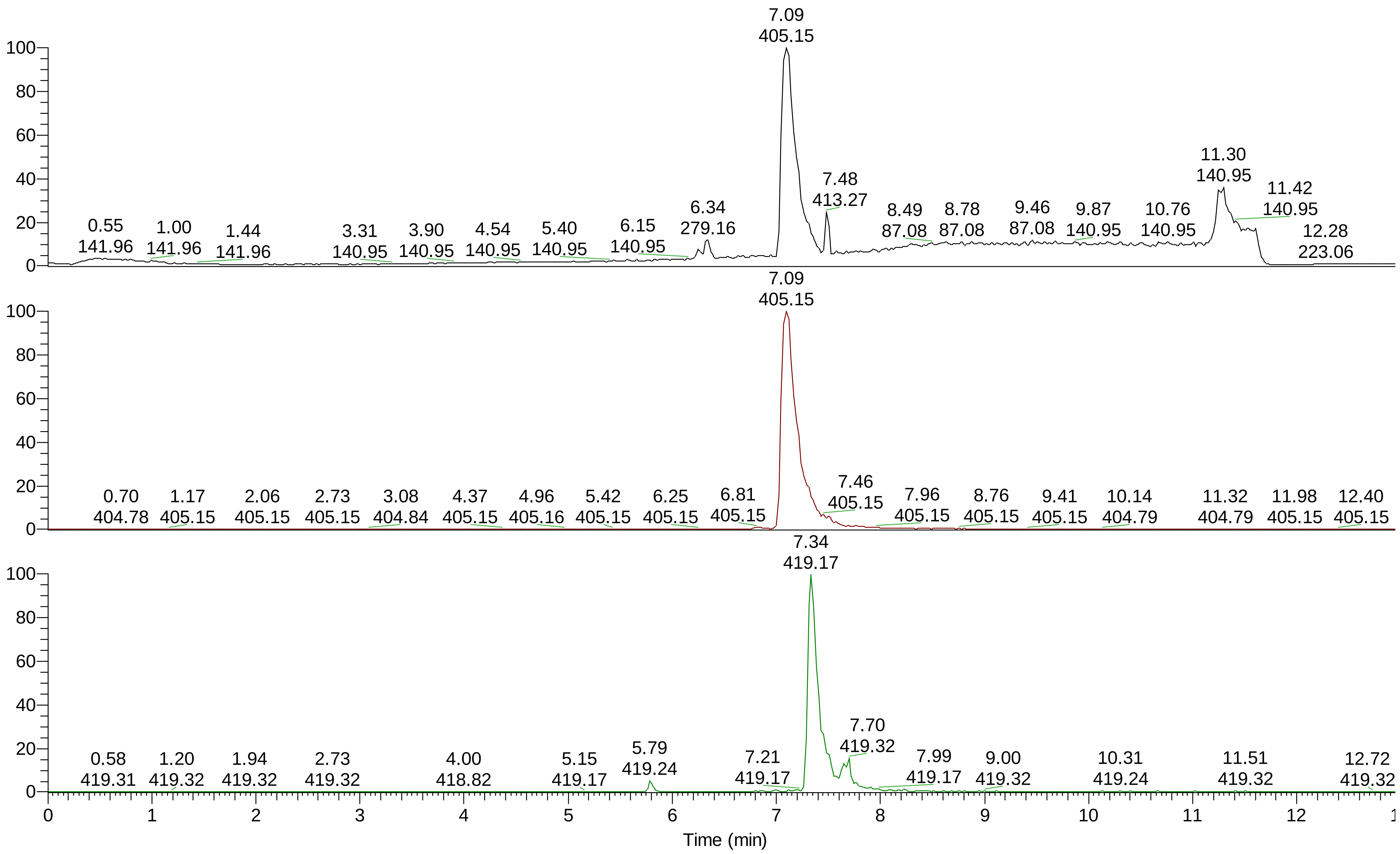


**Supplementary Figure 2: TIC chromatograms of HB5d/e3 (A3) at positive ionization.**

A: Total ion chromatograms of peak of HB5d/e3 (A3) at positive ionization (base peak intensity:1.86E8).. B: Mass range 404.5 - 405.5 was selected (base peak intensity: 3.63E8). C: Mass range 418.5 – 419.5.5 was selected (base peak intensity: 3.82E7).

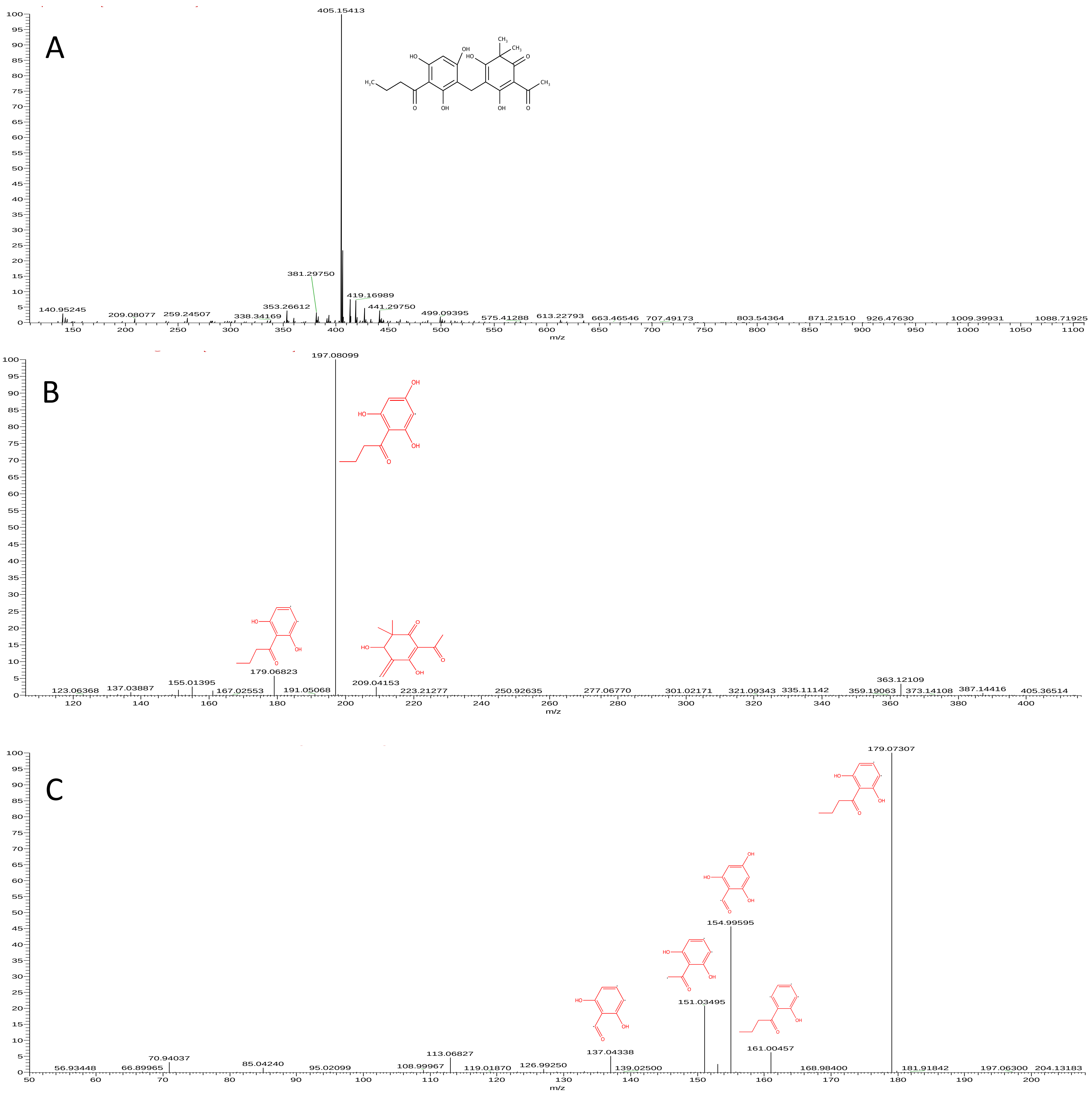


**Supplementary Figure 3: A: represents the conventional MS/MS spectra data of norflavaspidic acid AB ([MH]+obs. = 405.15413 amu).**

A peak at *m/z* 419 and 441 were contaminations from the column. B: Represents the spectrum of the MS/MS/MS experiment in which norflavaspidic acid AB was fragmented by in-source fragmentation and its product ion peak at *m/z* at 405 was selected by Q1 and further fragmented in Q2. A second-generation product ion at *m/z* 197 was a result of breaking of the bond. C: represents the MS/MS/MS spectrum of the product ion peak at *m/z* 197. A main product ion at m/z 179 accompanied a neutral loss of 18 amu. The spectra also contains other peaks at *m/z* 154(loss of loss of 25 amu), *m/z* 151 (loss of 28 amu), *m/z* 161(loss of 18 amu) and *m/z* 137 (loss of 42 amu).

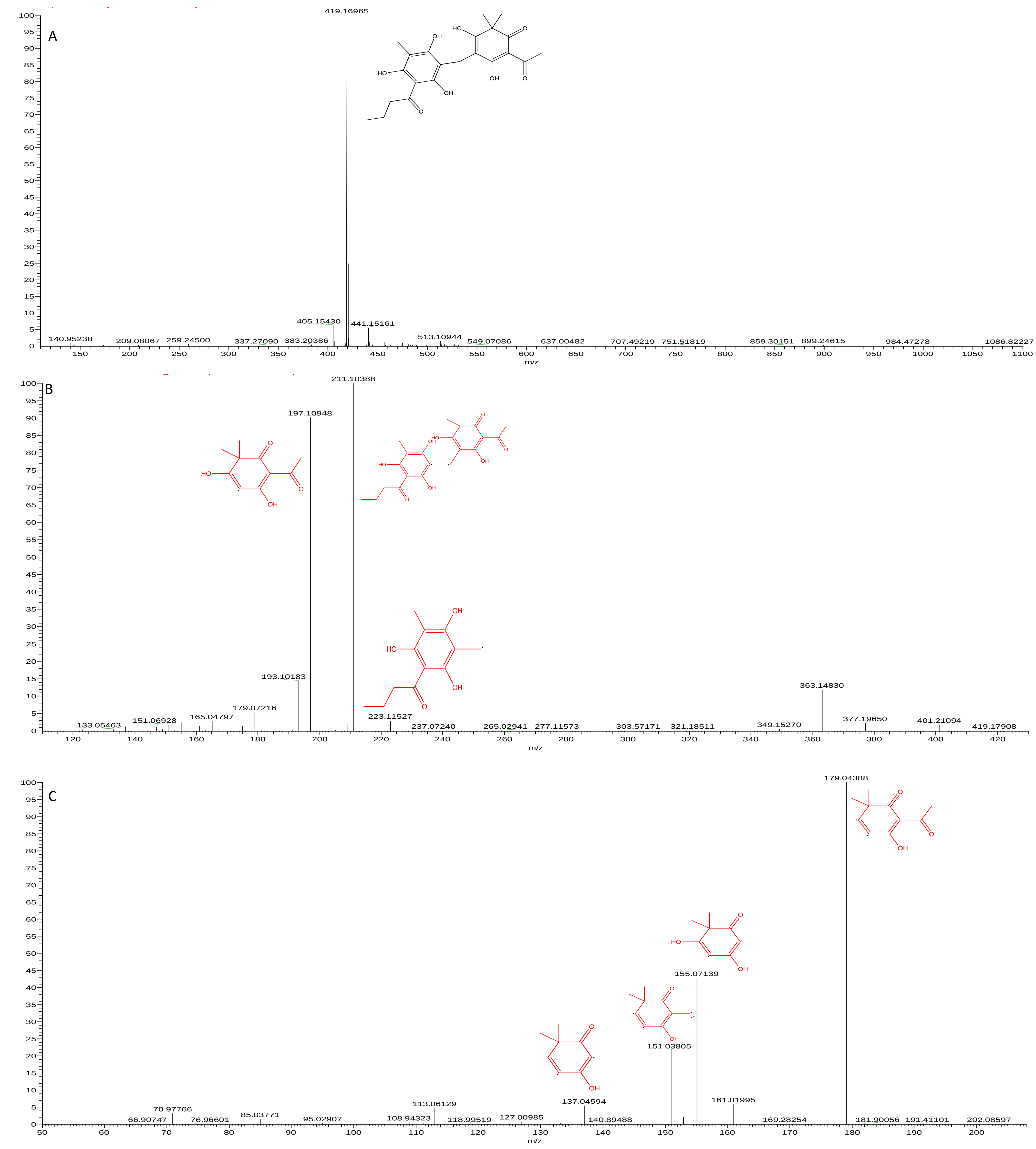


**Supplementary Figure 4: represents the conventional MS/MS spectra data of flavaspidic acid AB ([MH]+obs. = 405.16965 amu).**

A peak at *m/z* 441 were contaminations from the column. B: Represents the spectrum of the MS/MS/MS experiment in which flavaspidic acid AB was fragmented by in-source fragmentation and its product ion peak at *m/z* at 405 was selected by Q1 and further fragmented in Q2. A second-generation product ion at *m/z* 211 and 197 was a result of breaking of the bond. C: represents the MS/MS/MS spectrum of the product ion peak at *m/z* 197. A main product ion at *m/z* 179 accompanied a neutral loss of 18 amu. The spectra also contains other peaks at *m/z* 155(loss of loss of 24 amu), *m/z* 151 (loss of 28 amu), *m/z* 161(loss of 18 amu) and *m/z* 137 (loss of 42 amu).

**(A)**

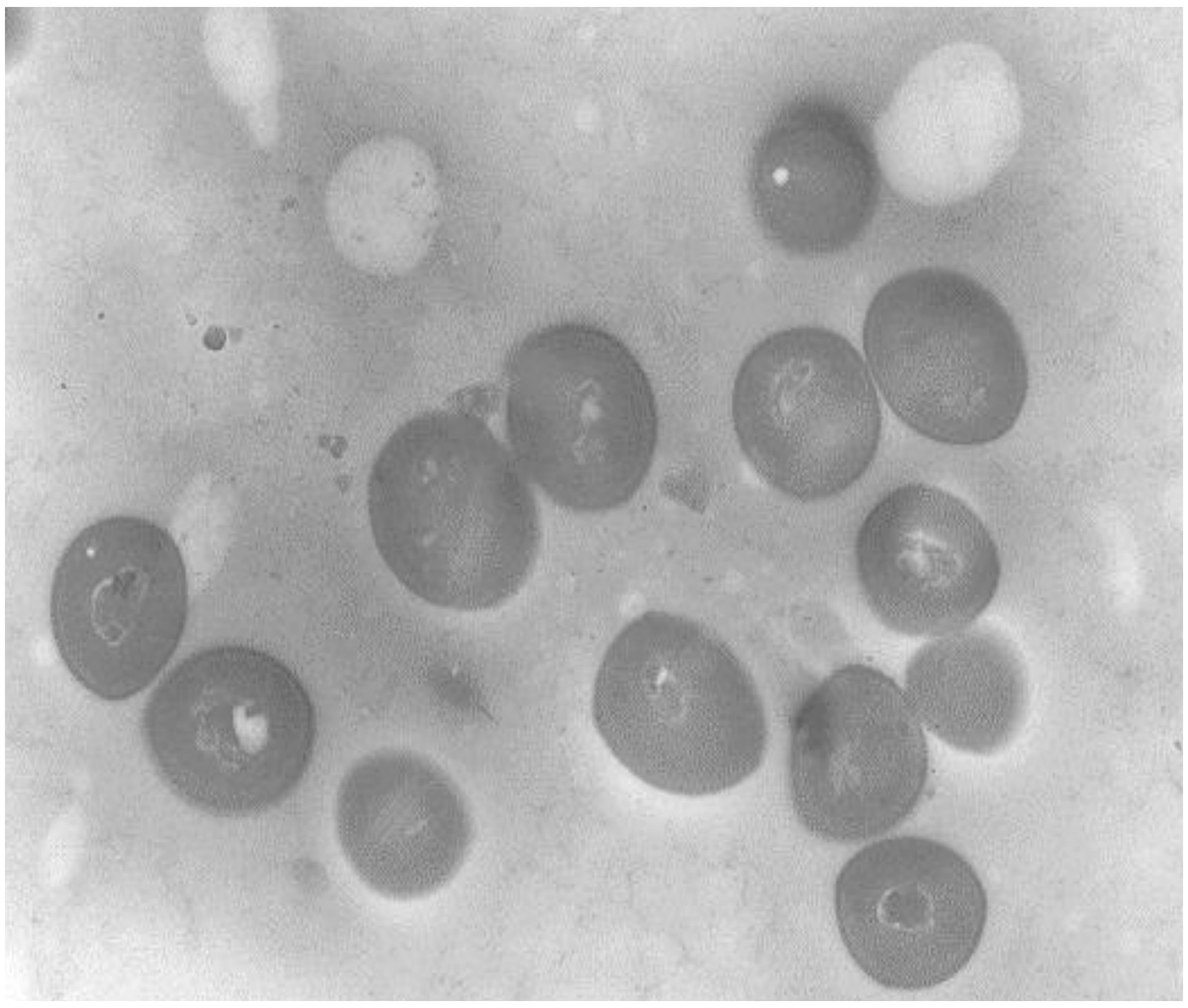

**(B)**

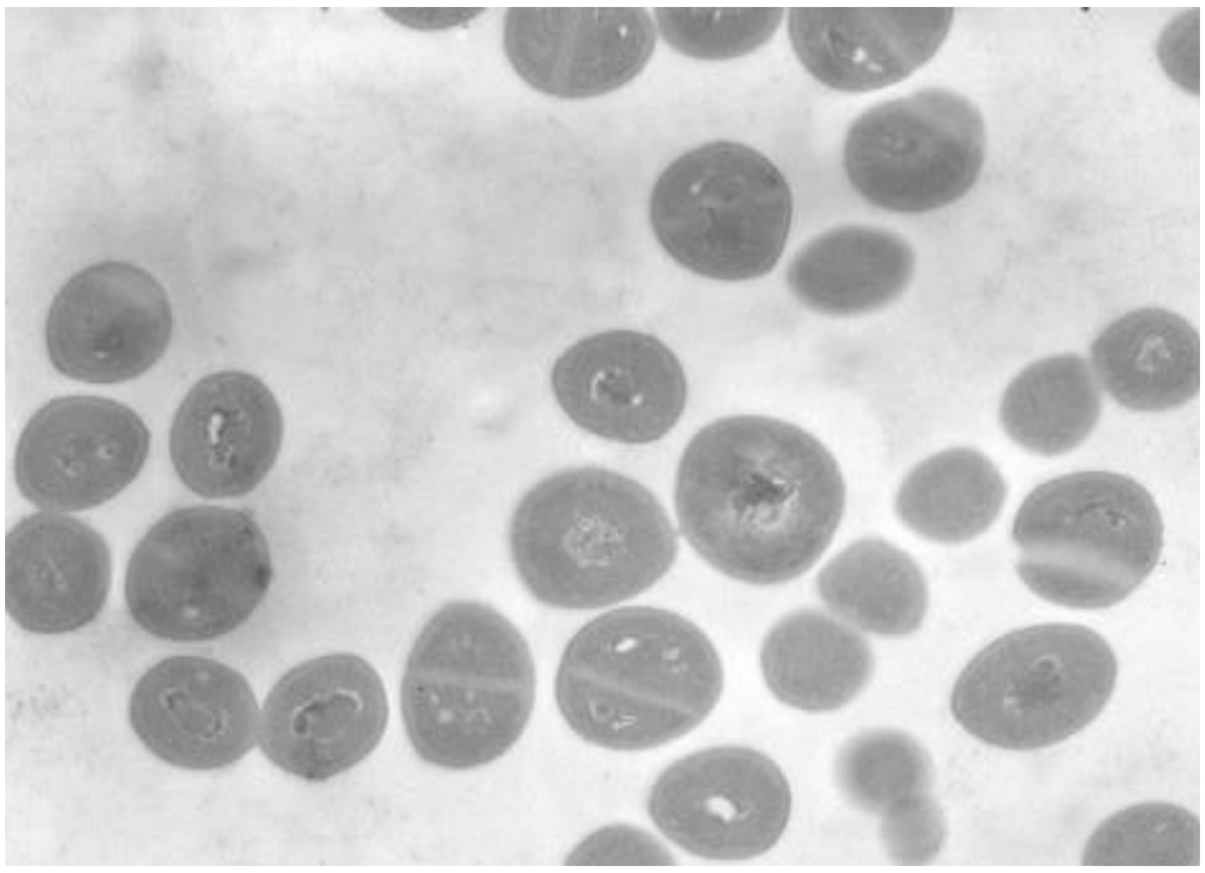

**(C)**

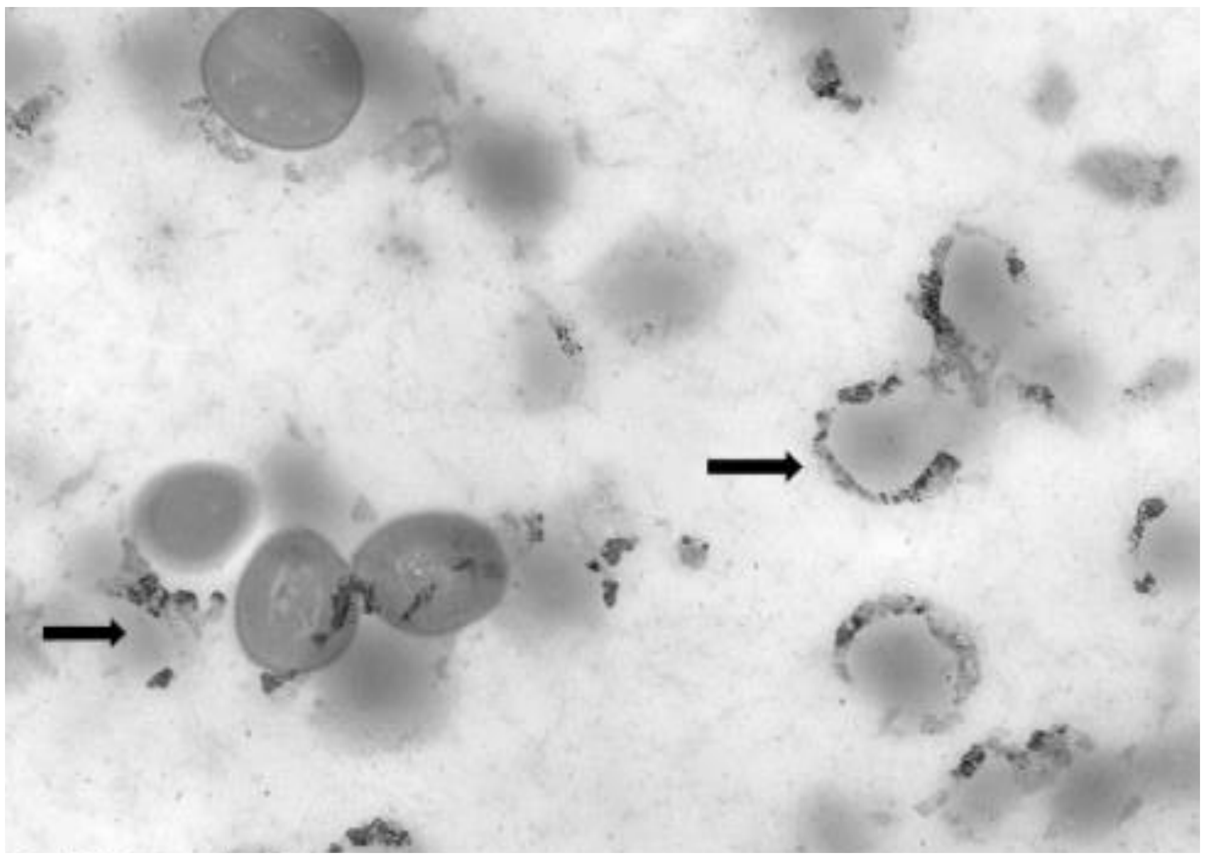

**Supplementary Figure 5 : Transmission electron microscope images of MRSA:USA 300 treated with two different concentration of HB5e**

(A) untreated controls at 6h (B) concentrations of HB5e sufficient to inhibit growth by 50 % at 6h ($MIC_{50}$) and (C), and contractions sufficient to inhibit growth by 90 % at $MIC_{90}$ at 24h. Arrow shows the cell membrane disruption and lysis

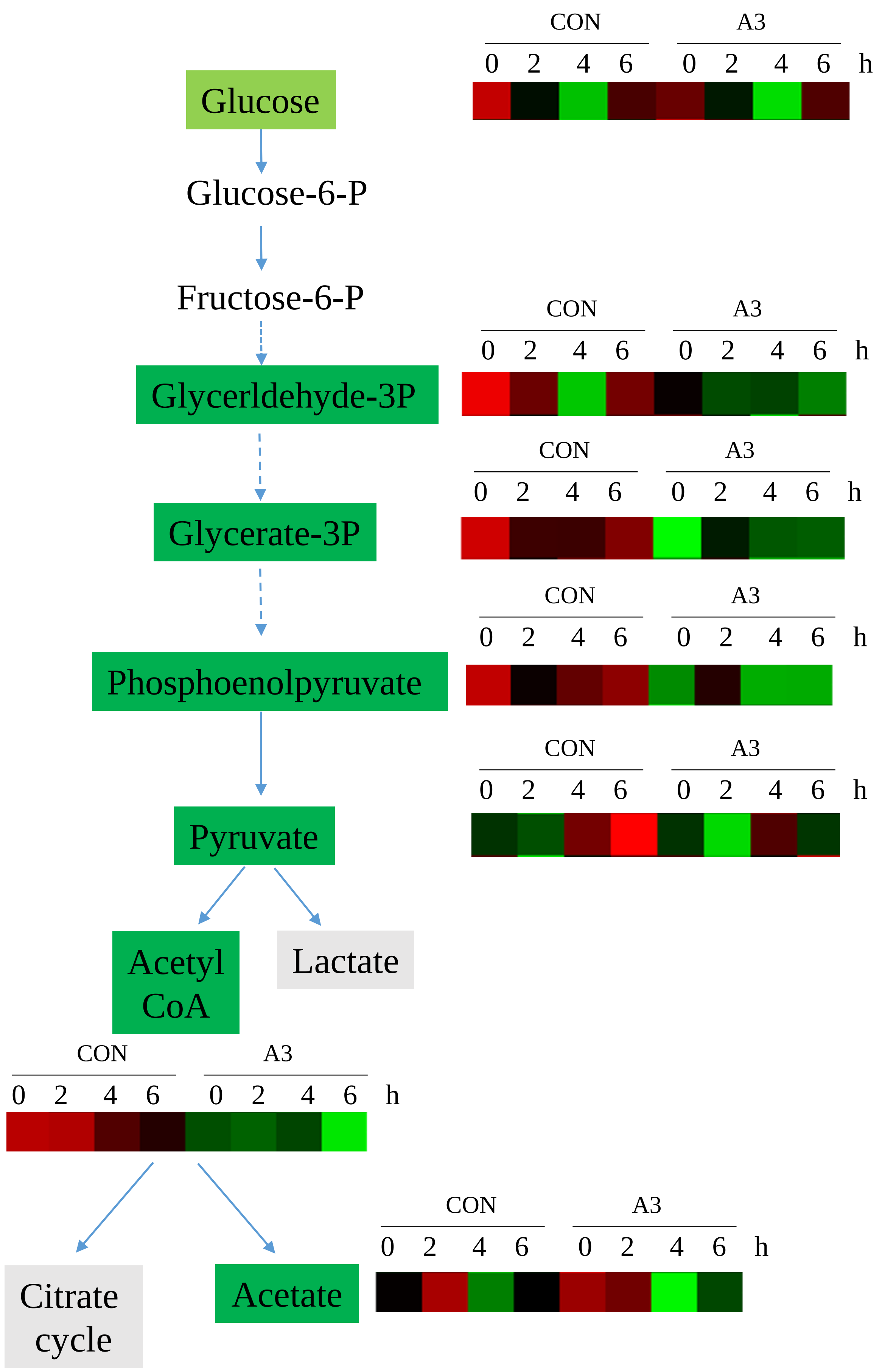


**Supplementary Figure 6 :** Schematic showing metabolite changes in MRSA cell following treatment with A3 based on KEGG map00010 "glycolysis and gluconeogenesis".

Metabolite names which were shown to be reduced with A3 treatment are indicated with a green background. The heatmap showing changes over time in controls (CON) and A3 treatment are also given (also provided in Figure 5). Metabolites or pathways highlighted in grey, showed no significant change in A3 treatment.
